# Interplay between abiotic conditions and mycorrhizal abundance determines differentiation and potential adaptation in a Mediterranean orchid

**DOI:** 10.64898/2026.02.25.707645

**Authors:** Marco G. Balducci, Karl J. Duffy

## Abstract

- Disentangling the relative influence of abiotic and biotic factors on plant population differentiation is a major challenge. Orchids often occur in patchily distributed populations, and all orchids depend on orchid mycorrhizal fungi (OrM) for seed germination. Hence, local abiotic conditions together with OrM may influence population differentiation and adaptation.
- Based on 316,952 polymorphic SNPs sampled from 21 populations throughout the range of the Mediterranean orchid, *Orchis italica* (Poir.), we performed a suite of analyses to test how population differentiation and potential adaptation is influenced by the interplay between abiotic factors and OrM.
- We found strong differentiation at the regional level, while loci under selection were associated with temperature, precipitation regime, soil texture, and overall OrM abundance. Outlier SNP functions were associated with stress responses and metabolic processes in the presence of OrM.
- Abiotic conditions and OrM combined determines differentiation in *O. italica*. Identifying selective pressures underlying differentiation and adaptive variation is critical for understanding plant responses to ongoing environmental change.

## Introduction

Genetic diversity is frequently partitioned among populations of species often leading to population differentiation and adaptation to local environments (Givnish, 2010). Abiotic factors, such as climate and soil conditions, play important roles in determining genetic differentiation of populations of species across large geographical ranges (Pearson & Dawson, 2003; Hawkins *et al*., 2003; Fischer *et al*., 2013), yet the role of biotic factors on genetic differentiation remains poorly understood (Baselga *et al*., 2012). Populations of most species encounter diverse abiotic and biotic conditions, yet not all species respond to environmental heterogeneity in the same way (Hartke *et al*., 2021; Gallegos, Hodgins, & Monro, 2023). For instance, species capable of long-distance dispersal often show weak signals of local adaptation and low genetic differentiation, while species with limited dispersal may adapt to local conditions and persist in environmentally heterogenous landscapes (Lenormand, 2002; Räsänen & Hendry, 2008). It is hence necessary to disentangle the relative contributions of abiotic and biotic variables to better understand how genetic structure and selection varies according to variation in their individual effects.

Genetic differentiation generally results as a consequence of gene flow between populations declining with increasing geographic distance, resulting in patterns of isolation by distance (IBD; Wright, 1943; Hoskin *et al*., 2005; Ramírez-Barrera *et al*., 2019). Under IBD, genetic drift is predicted to drive divergence, resulting in spatial dependency of genetic variation, where geographically proximate populations are more genetically similar that those further apart. However, populations may also be structured by environmental differences, leading to isolation by environment (IBE; Paz *et al*., 2015; Mosca *et al*., 2018). IBE arises when gene flow is constrained by natural selection across environmentally heterogenous landscapes (Wang & Bradburd, 2014; Toczydlowski & Waller, 2019). Mechanisms that result in IBE include selection against immigrants, reduced hybrid fitness, or biased rates of dispersal regulated by climatic conditions (Wright, 1943; Kawecki & Ebert, 2004; Wang & Bradburd, 2014). These processes can act independently or in concert to partition genetic variation among populations (Sexton, Hangartner, & Hoffmann, 2014). Whereas IBD results solely from distance, IBE may result from the combined influence of multiple abiotic and biotic factors, which in turn leads to local adaptation (Wang & Bradburd, 2014).

All of the estimated 28,000 orchid species (Chase *et al*., 2015; Christenhusz & Byng, 2016) produce dust-like seeds that lack endosperm, which likely represents an adaptation to wind dispersal (Arditti & Ghani, 2000; Otero & Flanagan, 2006; Hamrick & Trapnell, 2011). Orchid seeds therefore depend on fungi from a phylogenetically disparate guild, termed orchid mycorrhizal fungi (OrM), to germinate. Despite their capacity for long-distance dispersal, seed germination is often strongly spatially clustered (Jacquemyn *et al*., 2007). Orchids frequently form small, patchily distributed populations which could be due to preferences for particular habitats (i.e., abiotic factors) and/or by reliance on particular OrM for seed germination and establishment (McCormick & Jacquemyn, 2014; Duffy et al. 2019; McCormick, Whigham, & Canchani-Viruet, 2018; Phillips, Reiter, & Peakall, 2020). These ecological requirements combined have probably strongly influenced orchid distributions, yet the combined roles of abiotic and biotic factors in shaping differentiation in orchids remain poorly understood.

We investigated factors underlying population diversity and differentiation in the Mediterranean orchid, *Orchis italica* Poir. We used *de novo* double-digest RAD sequencing (ddRADseq) on individuals sampled throughout its Mediterranean range to; (1) determine patterns of fine- and broad-scale spatial genetic differentiation across populations and regions, (2) quantify how climatic and soil conditions, and a key biotic variable, OrM abundance, influence genetic differentiation, (3) identify candidate loci potentially under directional selection, and (4) correlate the signal of these candidate loci to both abiotic and biotic factors.

## Materials and methods

### Study species

*Orchis italica* is a diploid perennial orchid that occurs in small (10 -20 flowering individuals, occasionally to 200 flowering individuals) populations in the Mediterranean. Each plant produces a single terminal racemose inflorescence bearing between 10 and 50 pale pink to white flowers that open acropetally within a few days (Kretzschmar, Eccarius, & Dietrich, 2007). Flowering occurs from the beginning of March until the end of May. The species is self-compatible and is primarily pollinated by generalist bees, occasionally by butterflies and wasps (Pellegrino, Bellusci, & Musacchio, 2010; Pellegrino, Mahmoudi, & Palermo, 2021). While metabarcoding of OrM communities revealed a high diversity of fungal taxa, *in vitro* seed germination experiments showed that this species is primarily associated with the genus *Tulasnella*, particularly members of the *T. calospora* complex (Balducci, Calevo, & Duffy, 2025) and this association is maintained throughout adulthood (Balducci, Calevo, & Duffy, 2024).

### Sampling

Leaf tissue was collected from 21 populations of *O. italica* spanning its distribution, including 19 populations in southern Italy and two in southern Greece (Table S1). From each population, five adult plants were sampled, with individuals spaced at least 10 m apart to minimise the likelihood of sampling close relatives. In one Italian population (Potenza) and both Greek populations, only two individuals per population were sampled (Table S1). Leaves were dried in silica gel, stored at −20 °C, and later used for DNA extraction. Total genomic DNA was extracted from preserved leaf material using the DNeasy Plant mini kit (Qiagen, Hilden, Germany) according to the manufacturer’s protocol. DNA concentration and purity were verified using a NanoDrop 2000 Spectrophotometer (Thermo Fisher Scientific, Wilmington, DE, USA).

### OrM identification

For 17 populations, root samples were collected from the same adult individuals used for genomic analyses. The abundance and diversity of OrM associated with *O. italica* were previously described in Balducci *et al*. (2025). Briefly, total genomic DNA from highly colonised root fragments were extracted using the DNeasy Plant Mini kit (Qiagen, Hilden, Germany) following manufacture protocol. Amplification was performed using a semi-nested approach. The internal transcribed spacer (ITS) region was first amplified with three primer sets; (i) ITS1-OFa/ITS-OFb/ITS4-OF (Taylor & McCormick, 2008), (ii) ITS1/ITS4tul (Taylor & McCormick, 2008), and (iii) a pair specifically designed for Tulasnellaceae OrM fungi, 5.8S-OF/ITS4Tul (Vogt-Schilb *et al*., 2020). A second PCR step using the primer combination fITS9 (Ihrmark *et al*., 2012) and ITS4 (Tedersoo *et al*., 2014) were used to target the ITS2 region. Pooled amplicons were purified and sequenced on an Illumina NovaSeq platform (2 x 250 bp). Raw FASTQ files were processed in QIIME2 (Bolyen *et al*., 2019) using DADA2 (Callahan *et al*., 2016) for quality filtering and chimera removal, followed by operational taxonomic (OTU) clustering at 98% (USEARCH61; Edgar *et al*., 2011). Taxonomic classification was performed using the UNITE ITS database (version 8, 16.10.2022; (Abarenkov *et al*., 2010; Kõljalg *et al*., 2013).

### Double-digest RAD sequencing: library preparation and sequencing

Restriction-site associated DNA libraries were sequenced by IGA Technology Services (Udine, Italy) following their custom ddRADseq protocol, with minor modifications to Peterson et al. (2012). Enzyme combinations were chosen based on the estimated genome size of *O. italica* (8.7 Gbp; Bou Dagher-Kharrat et al., 2013). Genomic DNA was fluorometrically quantified, normalised to a uniform concentration, and 300 ng per sample was digested with 2.4 U each of *Sph*I and *Eco*RI (New England BioLabs) in a 30 μL reaction with CutSmart buffer, incubated at 37 °C for 90 min, followed by 65 °C for 20 min. Digested fragments were ligated to barcoded adapters (2.5 pmol) using 160 U of T4 DNA ligase (New England BioLabs) in a 50 μL reaction, incubated at 23 °C for 60 min, 20 °C for 60 min, and 65 °C for 20 min. Samples were multiplexed, purified with 1.5 volumes of AMPure XP beads (Agencourt), and size-selected on a BluePippin instrument (Sage Science) to retain fragments between 440–700 bp. Size-selected DNA was amplified using indexed primers and Phusion High-Fidelity PCR Master Mix (New England BioLabs) under the following thermal profile: 95 °C for 3 min; 10 cycles of 95 °C for 30 s, 60 °C for 30 s, and 72 °C for 45 s; and a final extension at 72 °C for 2 min. Amplified products were purified with AMPure XP beads (Beckman Coulter Inc.), and quantified with a Qubit 2.0 Fluorometer (Invitrogen) and Bioanalyzer DNA assay (Agilent Technologies). Sequencing was performed on an Illumina NovaSeq 6000 platform using 150 bp paired-end reads, according to the manufacturer’s instructions (Illumina, San Diego, CA).

### Quality filtering and variant calling

Raw reads were processed based on the Stacks software suite (Catchen et al., 2013). Demultiplexing of Illumina reads was performed with process_radtags, followed by read assembly for each sample with ustacks. A catalogue of loci was generated with cstacks, and individual samples were matched against this catalogue using sstacks and tsv2bam. The gstacks module was then used to assemble paired-end contigs, merge them with single-end loci, align reads to loci, and call SNPs. Loci were filtered with the ‘population’s program in Stacks v2.53 (Catchen et al., 2013), retaining only loci present in at least 75% of individuals across all samples (–R 0.75). To reduce the risk of paralogous sequences, loci with observed heterozygosity greater than 0.8 (--max-obs-het 0.8) were excluded.

### Genetic diversity and population differentiation

Genetic diversity and differentiation statistics were calculated using the ‘populations’ program in Stacks (Catchen et al., 2013). For each population, we estimated observed heterozygosity (H_o_), expected heterozygosity (H_e_), nucleotide diversity (π), and inbreeding coefficients (F_IS_). Pairwise population differentiation was assessed using F_ST_.

We inferred individual ancestry coefficients and evaluated population structure using the sparse non-negative matrix factorisation (sNMF) algorithm implemented in the R package ‘LEA’ v2.8.0 (Frichot et al., 2014; Frichot & François, 2015). Ten values of K (K =2-10) were tested with 20 iterations each, and the number of ancestral populations was evaluated using the cross-entropy criterion. Since structure-like methods may underestimate or misidentify K (Kalinowski, 2011; Lawson, Van Dorp, & Falush, 2018), we explored the full range of K values to assess potential hierarchical structure. We tested for population structure using discriminative analysis of principal components (DAPC) (Jombart, Devillard, & Balloux, 2010) in the R package ‘adegenet’. Cross-validation with 1,000 permutations was used to determine the number of principal components retained. K-means clustering solutions were compared using the Bayesian information criterion (BIC), and individuals were assigned to populations based on the retention of 60% of cumulative variance.

To complement clustering approaches, we reconstructed phylogenomic relationships using IQ-TREE v2.0.3 (Nguyen et al., 2015). Model selection was performed with ModelFinder Plus (-m MFP), including ascertainment bias correction (ASC; Lewis & Puterman, 2001). Nodal support was assessed with 1,000 ultrafast bootstrap replicates (-bb 1000). The resulting unrooted phylogeny was visualised and annotated using the Interactive Tree of Life (https://itol.embl.de/).

### Genetic variation associated with spatial, abiotic, and biotic factors

To test for IBD and IBE, we compared pairwise population genetic distances (Nei’s *D*) with geographic and environmental distances (see environmental data below) using Mantel tests. We used the “mantel” function from the R package “vegan” (Oksanen *et al*., 2019), with Spearman rank correlation and 9,999 permutations. Geographic distances among populations were calculated as Haversine distances using the “distm” function in the “geosphere” package (Hijmans *et al*., 2017). Environmental distances were calculated as Euclidean distances using the “dist” function.

Partial redundancy analysis (pRDA) was used to quantify the relative contributions of spatial distance, environmental predictors, and biotic interactions in explaining adaptive genetic variation across populations (Capblancq & Forester, 2021). This approach is well suited to genomic datasets as it does not rely on population grouping assumptions or equilibrium requirements inherent to FST-based analyses. Spatial autocorrelation was addressed by converting pairwise geographical distances to principal coordinates of neighbour matrices (PCNMs) using the ‘pcnm’ function, with significant spatial eigenvectors (PCNM 1 and PCNM 2) included as conditional variables. Environmental predictors included 19 WorldClim bioclimatic variables (1950-2000; Fick & Hijmans, 2017) and soil parameters from the LUCAS Database (ESDAC; Van Liedekerke, Jones, & Panagos, 2006; Ballabio *et al*., 2016), at 1 km resolution in Quantum GIS (QGIS Development Team, 2024). All variables were log-transformed to normalize scales. Forward selection (‘forward.sel’ function in ‘vegan’) with 9,999 permutations identified optimal predictors, while variance inflation factors (VIFs) using ‘vif.cca’ function controlled for multicollinearity among bioclimatic and soil variables. The ‘envfit’ function assessed predictor significance in explaining genetic variation.

For 17 of the 21 populations, we included abundance, as measured by the number of reads, of six OrM OTU found consistently detected in all adult individuals of *O. italica* (see Table S1; Balducci *et al*., 2025). Among these, tul26, tul27, tul28 (Tulasnellaceae) and cer15 (Ceratobasidiaceae) were the most abundant and consistently present OTUs detected across populations, making them ideal candidates for inclusion in explanatory models. Population allele frequencies were used as the response matrix, with three classes of predictors: (1) environmental, (2) geographic (PCNM as conditional variables) and (3) OrM abundances. Forward selection retained three key environmental variables: bio1 (annual mean temperature), bio16 (precipitation in the wettest quarter) and coarse fragments (which is a proxy measure of soil drainage). We implemented four pRDA models using the “rda” function in the “vegan” package (Oksanen *et al*., 2019): (1) abiotic-only (all 21 populations); and for the 17-population subset: (2) abiotic-only; (3) OrM fungi only; and (4) combined abiotic-biotic. Model significance, axis contributions and variable effects were assessed using permutation-based analysis of variance (ANOVA; 999 iterations).

### Putative loci under selection

To identify SNPs putatively under natural selection, we implemented a dual approach combining population differentiation tests and environmental association analyses (de Villemereuil *et al*., 2014). Differentiation based methods identify loci with greater genetic divergence among populations than expected under neutral evolutionary processes (e.g., gene flow, genetic drift). In parallel, environmental association analyses detect loci whose allele frequencies covary with environmental variables, potentially indicating local adaptation to specific climatic conditions (i.e., temperature and precipitation; Hoban *et al*., 2016). For population differentiation, we implemented two complementary methods: pcadapt (Luu, Bazin, & Blum, 2017), which uses principal component analysis to detect loci excessively correlated with population structure without requiring predefined populations; and OutFLANK (Whitlock & Lotterhos, 2015), which calculates FST based on predefined geographical regions (Table S1) while being robust to false positives. OutFLANK was run with default parameters and a q-value threshold of 0.1, corresponding to a 10 % false discovery rate (Benjamini & Hochberg, 1995). For environmental associations, we used pRDA on SNP loadings from the first three constrained axes (Forester et al., 2018). Candidate outliers were identified as SNPs exceeding 3.5 standard deviations from the mean loading (p < 0.0005). When loci were associated with multiple RDA axes, Pearson’s correlation was used to assign the strongest predictor.

To provide functional context, candidate SNPs identified by pcadapt and pRDA were mapped to the *Platanthera zijinensis* reference genome (Li et al., 2022), an annotated genome available for a terrestrial orchid within the Orchidoideae. Genomic coordinates of outlier SNPs were intersected with genome annotations using BEDTools v2.30.0 (Quinlan, 2014) to identify overlapping or nearest genes. Functional information (gene ID, gene name, and predicted product) was extracted from the reference genome annotation, and genes were grouped into broad functional categories based on annotated product descriptions. Although reference-guided filtering reduced the number of retained SNPs relative to *de novo* analyses, this approach enabled biologically meaningful interpretation of candidate loci and allowed genomic signals of selection to be linked to functional pathways relevant to orchid biology.

## Results

### Population diversity and differentiation

After quality filtering to remove loci with high missing data, excessive heterozygosity, and low coverage, we retained 316,952 genotyped SNPs from 96 individuals representing 21 populations (Table S1). Italian populations (excluding Anacapri) showed consistently high levels of genetic diversity, with nucleotide diversity (π) ranging from 0.051 to 0.076, observed heterozygosity (H_o_) between 0.054 and 0.068, and expected heterozygosity (H_e_) from 0.032 to 0.067 (Table S1). In contrast, the two Greek populations and Anacapri showed significantly reduced diversity across all metrics (p < 0.005, Table S1). Private alleles were detected in all populations, ranging from 53 in Hymettus (Greece) to 14,580 in Acate (Sicily). The inbreeding coefficient (F_IS_) was positive in 17 of 21 populations, indicating widespread inbreeding. Pairwise F_ST_ values ranged from 0.064 to 1 (Fig. S1), and all comparisons were significantly greater than zero. The strongest differentiation was observed in the Greek populations, which were completely isolated from Italian populations (F_ST_ = 1).

### Association of genomic variation with geography and environment

There was a positive correlation between geographic and genetic distances, supporting the IBD hypothesis (r = 0.347, p = 0.01; Fig. S2). In contrast, based on the raw genetic distance matrix, the IBE hypothesis was not supported, with no association between environmental and genetic distances (r = 0.239, p = 0.407; Fig. S3).

However, partial RDAs controlling for geographic distance revealed that both abiotic and OrM communities significantly influenced genetic structure. Across the full dataset, abiotic predictors explained a significant proportion of genetic variation (p = 0.001, Fig. 1a), while in the 17-population subset the effect was weaker, yet still significant (p = 0.045, Fig. 1b). OrM community composition alone had a strong effect (p = 0.001, Fig. 1c), with the combined abiotic + OrM model providing the best fit overall (p = 0.005, Fig. 1d).

**Fig. 1.**
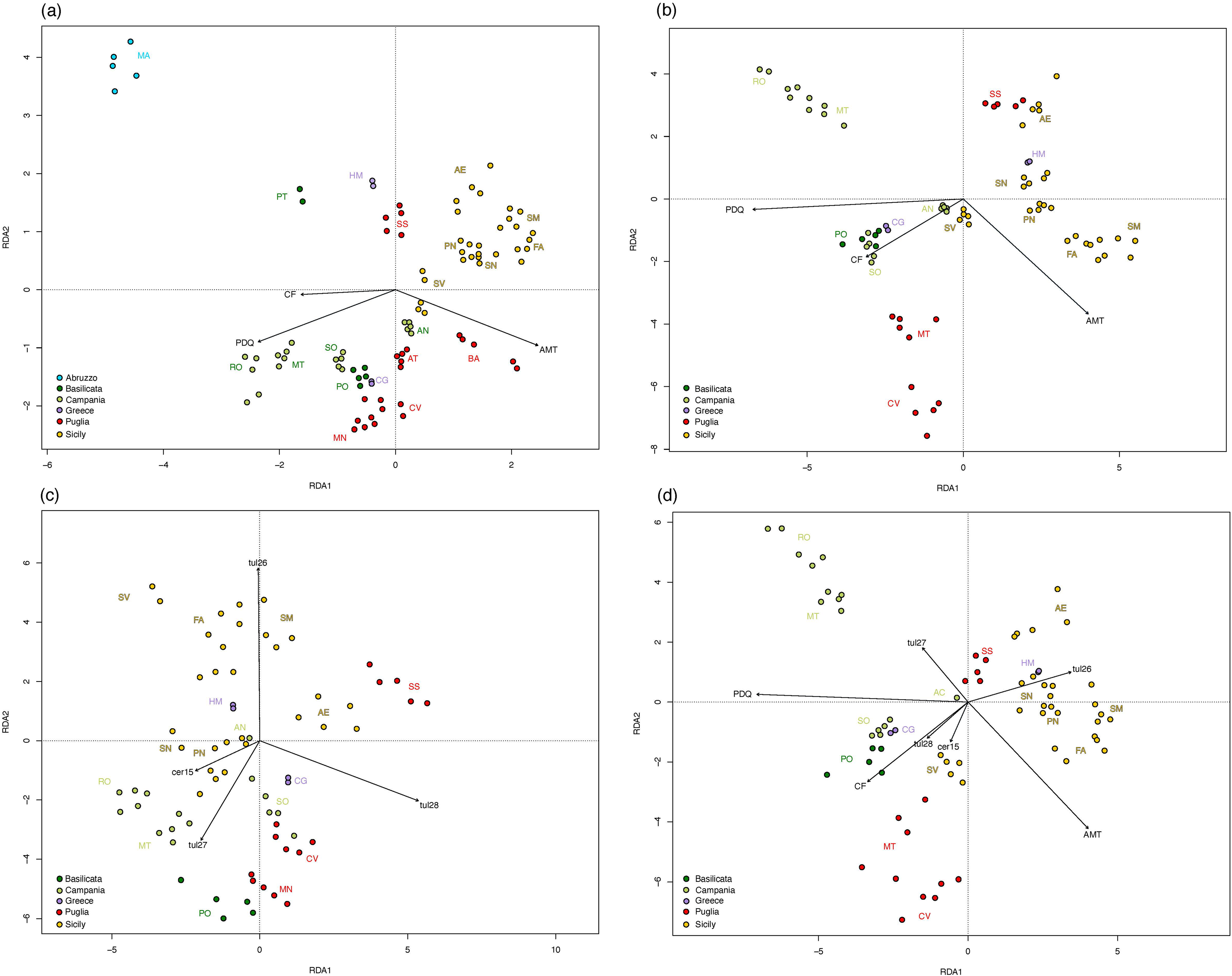
Partial RDAs of *O. italica* allele frequencies, showing the effects of coarse fragments (CF), precipitation of the driest quarter (PDQ) and annual mean temperature (AMT) on; (a) all populations, (b) the subset of 17 populations, (c) mycorrhizal communities, and (d) combined factors on genetic structure.

The full population analysis showed distinct genetic clustering in central and southern Italy (particularly Majella), with individuals from Roccamonfina and Matese diverging along a precipitation gradient (Fig. 1a). In the 17-population subset, genetic variation in Pollino (Basilicata), Hymettus (Greece), and Sorrento (Campania) was strongly associated with soil drainage (coarse fragments; Fig. 1b). In Sicily, populations from Santa Caterina Villarmosa, Ferla, and Santa Maria were positively associated with tul26, while Santa Ninfa and Partanna showed negative associations with cer15 (Fig. 1c). Northern Italian populations, along with Roccamonfina and Matese, were negatively associated with tul27 (Fig. 1c). The combined model explained the greatest variation, identifying tul26 as the primary driver for Sicilian populations, coarse fragments for central Italian populations (Pollino and Sorrento) and cer15 for Santa Caterina Villarmosa (Sicily; Fig. 1d).

We identified a total of 515 SNP outliers across all analyses (complete dataset in Table S2), including 115 from abiotic-only associations, 116 from OrM-only associations, and 47 from combined abiotic-biotic models. For all 21 populations, six candidate loci were associated with precipitation of the driest quarter, while 279 and 230 showed significant associations with coarse fragments and annual mean temperature, respectively (Fig. S4a). Forty-one candidate outliers overlapped with those detected by pcadapt. In the 17-population subset with OrM data, annual mean temperature showed the strongest association (68 SNPs), followed by precipitation of the driest quarter (37 SNPs) and coarse fragments (10 SNPs) (Fig. S4b). OrM-specific analyses revealed 107 SNPs linked to tul28, with fewer associations for tul27 (7 SNPs), tul26 (1 SNP), and cer15 (1 SNP) (Fig. S4c). The combined model identified loci associated with precipitation of the driest quarter (36 SNPs), coarse fragments (5 SNPs), tul28 (5 SNPs), and tul27 (1 SNP) (Fig. S4d).

### Population structure

(DAPC identified K = 7 as the most likely number of clusters based on the BIC and find.clusters() function. Populations generally grouped together irrespective of geographic distance, except for Majella (Abruzzo, central Italy), which formed a distinct cluster (Fig. S5). sNMF analysis indicated an optimal K = 5, but closer inspection showed that K = 3 best captured the major genetic clusters in *O. italica*, as higher values of K reflected substructure rather than geographically meaningful population groups. Admixture analysis at K = 3 revealed three primary ancestral components: (1) Majella, (2) Sicily, and (3) mainland Italian and Greek populations (Fig. 2a). Additional substructure emerged at higher K values. Maximum likelihood phylogenetic analysis, using the best-fit K3Pu + F + R6 model (selected by BIC), recovered strong geographic clustering by region, consistent with admixture results (Fig. 2b). Principal component analysis based on putative outlier SNPs further supported this structure, with populations separating along the first two axes consistent with regional clustering, particularly distinguishing Abruzzo and Sicilian populations (Fig. 2c).

**Fig. 2.**
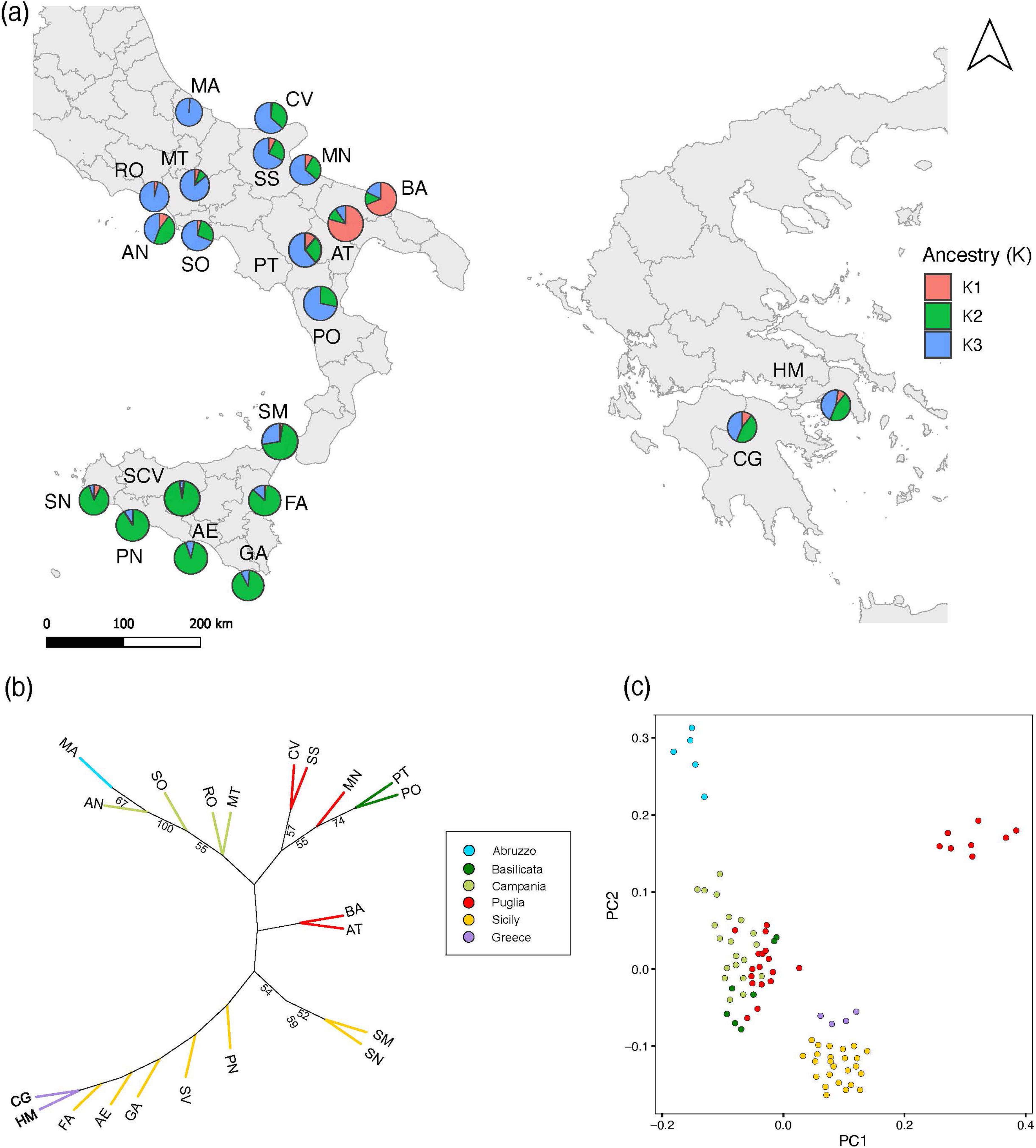
(a) Admixture plots obtained from sNMF analysis showing hierarchical population structure of *Orchis italica* across different K values. Population names are abbreviated as in Table S1. (b) Unrooted maximum likelihood tree for 21 populations of *O. italica* from southern Italy and Greece. Each tip represents an individual population which is colour coded to its region. Branch labels indicate support from 10,000 ultrafast bootstrap replicates. (c) Genetic variation based on putative outlier SNPs among populations of *O. italica* along the first two axes of a principal component analysis.

### Putative loci under selection

Mapping candidate SNPs to the *P. zijinensis* reference genome revealed that 234 outlier SNPs identified across all analyses could be confidently placed within annotated genomic regions (Fig. S6). Of these, 65 SNPs were detected by pcadapt (Fig. 6a), while 187 SNPs were identified by environmental association analyses, including 100 associated with OrM abundance (Fig. 6b), 120 associated with abiotic variables (Fig. 6c), and 127 detected in the combined abiotic-biotic model (Fig. 6d), with partial overlap among models. Eighteen SNPs were shared between pcadapt and RDA approaches, indicating largely complementary signals of selection (Table S3).

Most mapped outlier SNPs were located within or near annotated genes, frequently within coding sequences or untranslated regions. Functional annotation revealed that genes linked to candidate loci were enriched for roles in signal transduction, stress and defence responses, transcriptional regulation, and metabolism. Recurrently detected gene classes included receptor-like kinases, transcription factors, oxidoreductases, and enzymes involved in primary metabolism, consistent with genomic responses to environmental and biotic variation. Notably, SNPs identified exclusively by the biotic model were often associated with genes involved in cell signalling, redox regulation, and metabolic control. Many of these loci were absent from abiotic-only models, suggesting that OrM associations impose selective pressures not captured by climatic or edaphic variables alone. A large proportion of candidate SNPs, however, mapped to genes of unknown or hypothetical function, reflecting both incomplete annotation of orchid genomes and the likelihood that additional, lineage-specific mechanisms of adaptive divergence remain uncharacterised.

## Discussion

The interplay between abiotic and biotic variables influences population differentiation and potential adaptation to local environments. Our results indicate that the Mediterranean orchid, *O. italica*, has pronounced genetic differentiation, which likely reflects a combination of historical isolation and contemporary barriers to gene flow. We found that Greek populations were genetically distinct from Italian populations. Similar disjunct patterns may be common in the Mediterranean flora, where geographic discontinuities correspond to strong phylogeographic structuring (Médail & Diadema, 2009; Nieto Feliner, 2014). While most Italian populations have relatively high genetic diversity (as estimated by H_o_, H_e_, and *π*), the reduced diversity in the Greek populations suggests prolonged geographic and reproductive isolation. Comparable reductions in diversity across Mediterranean orchid populations have been reported in other orchid species, such as *Dactylorhiza insularis* (Bullini *et al*., 2001). Genetic diversity can be similar from populations sampled at both the limits and centre of a species range (see for example Duffy et al., 2009), yet even at very local scales, genetic diversity can shift according to population (Duffy et al., 2020).

The distribution of private alleles provided further insight into differentiation. Eastern Sicilian populations (Acate, Ferla and Gela) harboured elevated numbers of private alleles, consistent with long-term isolation that restricted gene flow and increased genetic drift. These alleles may represent retained ancestral variants persisting from formerly fragmented refuge populations, or they may reflect locally adapted polymorphisms shaped by unique environmental conditions. Such signals of local differentiation are common in Mediterranean orchids, where island and peninsular populations often exhibit high levels of genetic distinctiveness (Cozzolino & Widmer, 2005; Tremblay et al., 2005). In contrast, the two Greek populations (Hymettus and Mount Kyllini), and the Italian island population of Anacapri contained relatively few private alleles, suggesting recent colonisation events, limited time for divergence, or ongoing gene flow that homogenises genetic diversity across sites.

Despite regional variation in diversity and allele composition we found consistently low F_IS_ values, yet F_IS_ generally increased with geographic distance. This lack of strong inbreeding is likely maintained by a combination of life-history traits. First, strong selection against inbred progeny during early developmental stages reduces the persistence of selfed lineages. Second, as *terrestrial orchids are typically long-lived perennials with* extended generation overlap (Tremblay et al., 2005), demographic buffering may reduce the stochastic effects of drift. Third, pollinator-mediated outcrossing due to deceptive pollination characteristic of *Orchis*, where pollinator learning reduces within-population revisitation while enhancing pollen transfer among populations (Neiland & Wilcock, 1998; Cozzolino & Widmer, 2005; Minasiewicz et al., 2018). Together, these features contribute to the maintenance of genetic variation despite the fragmented and patchy distribution typical of Mediterranean orchid populations.

Our analyses revealed that geographically proximate populations of *O. italica* are genetically similar, consistent with IBD. This indicates that genome-wide neutral genetic structure is primarily shaped by geographic distance. This pattern mirrors findings from a wide range of plant taxa, in which spatial separation and physical barriers dominate genome-wide differentiation, even across environmentally heterogenous landscapes (Holderegger et al., 2010; Sexton et al., 2014; Mosca et al., 2018). Limited dispersal and historical connectivity structures neutral variation, while environmental gradients primarily affect a subset of loci under selection. Our pRDA analyses indicate that particularly in combination with OrM OTU read abundance, climate (temperature and precipitation), and soil properties are putative drivers of adaptive differentiation. Comparable patterns have been observed in alpine and Mediterranean plants, where IBD governs neutral background structure but local adaptation to fine-scale environmental heterogeneity drives selection on specific genomic regions (Frei et al., 2012; De Kort et al., 2014; Capblancq et al., 2020). This distinction between neutral and selective processes has important implications for predicting the capacity of *O. italica* to persist under climate change. Neutral genetic differentiation shaped by IBD suggests that dispersal limitation will constrain natural range shifts, as seen in other orchids with restricted seedling recruitment away from adult populations (Jacquemyn et al., 2012). At the same time, the environmental associations of candidate loci imply that local adaptation may depend on narrow climatic thresholds and soil properties, mirroring findings in other species where adaptive genomic variation tracks microenvironmental gradients (Jia et al., 2020; Gugger et al., 2021). Together, these results highlight the dual role of geography and environment in shaping the evolutionary trajectory of *O. italica* – geographic distance restricts neutral connectivity, while selection linked to environmental heterogeneity and mutualists (OrM) drive adaptive divergence.

Differentiation was strongly associated with abiotic variables such as temperature, precipitation, and soil conditions. The Mediterranean is characterised by dry, hot summers and cold, comparatively wet winters (Cowling *et al*., 2015). Precipitation regimes may impose strong constraints on plants and can act as an important evolutionary driver for adaptation to both drought conditions in the summer and inundation in the winter (Galmés, Medrano, & Flexas, 2007). For *O. italica*, plants typically grow in open grasslands on well-drained calcareous soils, hence summer droughts, which coincide with early onset of seed germination and development, may impact recruitment. Precipitation regimes, along with higher summer temperatures, may also shape the composition of OrM communities associated with *O. italica* (Balducci *et al*., 2025).

Our results provide genome-scale evidence that OrM may contribute directly to local adaptation. By integrating SNP markers with fungal abundance patterns, we showed that specific OrM taxa are linked to genetic structure in *O. italica*, potentially acting as biotic filters that influence patterns of adaptation. This was particularly evident in the Sicilian populations, where strong associations with tul26 and cer15 suggest that these OrM OTUs exert variable selective pressures. Such patterns imply that OrM associations can mediate local adaptation to particular ecological conditions, consistent with findings from other orchids where shifts in fungal partners correspond to population differentiation and recruitment success (Jacquemyn et al., 2011; Těšitelová et al., 2013). Our results therefore suggest that OrM are not mere passive facilitators of orchid seed germination, and highlight that they may act as active drivers of divergence. Indeed, SNPs detected exclusively by the OrM-only pRDA were associated with genes involved in cell signalling, redox regulation, metabolism, and transcriptional control, functions consistent with plant-fungal interactions and symbiotic nutrient exchange (Parniske, 2008; Bonfante & Genre, 2010; Smith & Read, 2010). The absence of these loci from abiotic-only models indicates that mycorrhizal fungi impose selective pressures partly independent of climatic and edaphic gradients, acting as biotic filters shaping host genomic variation. Genes linked to redox balance and metabolic regulation are particularly relevant given their roles in carbon-nutrient exchange and oxidative signalling during symbiosis (Bago et al., 2000; Genre et al., 2012), while the recurrent detection of signalling-related genes and transcriptional regulators reflects the molecular coordination required to maintain mutualistic balance (Kiers et al., 2011; Wang et al., 2021). Together, these patterns link variation in OrM abundance to divergence at functionally relevant host loci, supporting a role for biotic interactions in adaptive divergence alongside abiotic drivers.

Population structure analyses further supported these patterns. Both DAPC and sNMF revealed consistent clustering, with individuals from Abruzzo forming a distinct genetic group. This pronounced differentiation likely reflects the region’s steep environmental gradients and mountainous topography, which may act as barriers to gene flow by limiting both pollinator movement and seed dispersal (Steinbauer et al., 2016; Wu et al., 2020). Sicilian populations also showed a distinct genetic profile, although weaker than the divergence observed in Abruzzo. In contrast, high connectivity was observed among populations from Basilicata, Campania, and Puglia, suggesting either historical gene flow or ongoing long-distance dispersal. Nevertheless, even within these well-connected regions, we detected localised differentiation, such as between the geographically proximate populations of Altamura and Bari in Puglia. This fine-scale structure may reflect independent colonisation events from distinct founder populations or subsequent reproductive isolation despite close geographic proximity.

## Conclusions

While geographic distance between populations primarily structures neutral genomic variation, differentiation is strongly influenced by the combination of abiotic variables and abundance of OrM fungi. Such mutualistic fungi may act as selective agents shaping population structure in *O. italica*, highlighting the broader role of biotic interactions in plant evolution. Hence, adaptation is not solely a function of climate and soil but also depends on the ecological distribution of compatible fungal partners. Recognising OrM as potential co-drivers of local adaptation should focus orchid conservation strategies on protecting not just plants but also the belowground ecological networks that sustain them.

## Acknowledgments

We are grateful to Jacopo Calevo for assistance with leaf sample collection and assistance in the field, and Spyridon Oikonomidis for contributing samples from Greece. We gratefully acknowledge funding from the Italian Ministry of Research (PRIN grant number 2017TP3SJL, awarded to KJD).

## Data Availability Statement

Raw ddRAD sequencing data have been deposited in the NCBI Sequence Read Archive under BioProject accession PRJNA1427967. Individual run accessions range from SRR37357204–SRR37357298. Processed data files will be archived in Dryad upon manuscript acceptance.

## Supporting information

Additional supporting information may be found online in the Supporting Information section at the end of the article.

---

**Fig. S1.** Heat map of pairwise F_ST_ values for 21 populations of *O. italica*. Deeper colours indicate higher genome-wide differentiation between populations, according to the colour key. Population names are abbreviated as in Table S1.

**Fig. S2.** Positive isolation}-by-distance relationship (Mantel test: r = 0.347, p = 0.01) between genetic and geographic distance across populations of *Orchis italica*.

**Fig. S3.** Relationship between genetic distance and environmental distance across populations *Orchis italica*.

**Fig. S4.** Partial RDAs of *O*. *italica* allele frequencies, showing candidate SNPs coloured by their most strongly correlated abiotic and biotic variables. Panels show RDAs for: (a) all 21 populations, and (b) a subset of 17 populations. Variables include coarse fragments (CF), precipitation of the driest quarter (PDQ), and annual mean temperature (AMT). Panels (c) and (d) show the effects of (c) mycorrhizal communities and (d) combined abiotic and biotic factors on genetic structure in the 17-population subset.

**Fig. S5.** Scatterplots resulting from discriminant analysis of principal components without prior population information.

**Fig. S6.** Distribution of functional gene categories associated with candidate adaptive SNPs. Counts represent genes linked to outlier SNPs detected by pcadapt and partial redundancy analyses (abiotic, biotic, and combined models). SNPs were annotated using the *Platanthera zijinensis* genome and grouped into broad functional classes based on predicted gene function.

## Tables

Table S1. Summary genetic statistics for all populations for only variant nucleotide positions that are polymorphic in at least one individual. These statistics include the number of leaves and roots collected from each population, the number of variable sites unique to each population (private), the average observed heterozygosity per locus (H_O_), the average expected heterozygosity per locus (H_E_) the average nucleotide diversity (π), and the average Wright’s inbreeding coefficient (F_IS_).

Table S2. Outlier SNP correlations with environmental predictors: AMT (Annual Mean Temperature), PDQ (Precipitation Driest Quarter), CF (Coarse Fragments), and mycorrhizal fungi (cer15, tul26, tul27 and tul28). Values show correlation strength by model type (abiotic, biotic, or combined).

Table S3. Functional annotation of candidate outlier SNPs associated with abiotic and biotic predictors in *Orchis italica*. SNPs identified by pcadapt and partial redundancy analysis (pRDA) are mapped to the *Platanthera zijinensis* reference genome. For each SNP, the table reports genomic position, associated environmental or mycorrhizal predictor(s), nearest or overlapping gene, gene annotation, and predicted functional category. Overlap among detection methods is indicated where applicable.

## References

Abarenkov K, Nilsson RH, Larsson KH, Alexander IJ, Eberhardt U, Erland S, Høiland K, Kjøller R, Larsson E, Pennanen T. 2010. The UNITE database for molecular identification of fungi–recent updates and future perspectives. The New Phytologist 186: 281–285.

Arditti J, Ghani AKA. 2000. Tansley Review No. 110. Numerical and physical properties of orchid seeds and their biological implications. New Phytologist 145: 367–421.

Bago B, Pfeffer PE, Shachar-Hill Y. 2000. Carbon metabolism and transport in arbuscular mycorrhizas. Plant Physiology 124: 949–958.

Balducci MG, Calevo J, Duffy KJ. 2025. Orchid mycorrhizal communities associated with *Orchis italica* are shaped by ecological factors and geographical gradients. Journal of Biogeography 52: 544–557.

Balducci MG, Calevo J, Duffy KJ. 2024. Narrow mycorrhizal specialization and its effect on early development in the Mediterranean orchid *Orchis italica*. Botanical Journal of the Linnean Society 206: boae018.

Ballabio C, Panagos P, Monatanarella L. 2016. Mapping topsoil physical properties at European scale using the LUCAS database. Geoderma 261: 110–123.

Baselga A, Lobo JM, Svenning J, Araújo MB. 2012. Global patterns in the shape of species geographical ranges reveal range determinants. Journal of Biogeography 39: 760–771.

Benjamini Y, Hochberg Y 1995. Controlling the false discovery rate: a practical and powerful approach to multiple testing. Journal of the Royal Statistical Society: Series B (Methodological) 57: 289–300.

Bolyen E, Rideout JR, Dillon MR, Bokulich NA, Abnet CC, Al-Ghalith GA, Alexander H, Alm EJ, Arumugam M, Asnicar F. 2019. Reproducible, interactive, scalable and extensible microbiome data science using QIIME 2. Nature Biotechnology 37: 852–857.

Bonfante, P, Genre A. 2010. Mechanisms underlying beneficial plant–fungus interactions in mycorrhizal symbiosis. Nature Communications 1: 1–11.

Bou Dagher-Kharrat M, Abdel-Samad N, Douaihy B, Bourge M, Fridlender A, Siljak-Yakovlev S, Brown SC. 2013. Nuclear DNA C-values for biodiversity screening: Case of the Lebanese flora. Plant Biosystems 147: 1228–1237.

Bullini L, Cianchi R, Arduino P, De Bonis L, Mosco MC, Verardi A, Porretta D, Corrias B, Rossi W. 2001. Molecular evidence for allopolyploid speciation and a single origin of the western Mediterranean orchid *Dactylorhiza insularis* (Orchidaceae). Biological Journal of the Linnean Society 72: 193–201.

Callahan BJ, McMurdie PJ, Rosen MJ, Han AW, Johnson AJA, Holmes SP. 2016. DADA2: High-resolution sample inference from Illumina amplicon data. Nature Methods 13: 581–583.

Capblancq T, Morin X, Gueguen M, Renaud J, Lobreaux S, Bazin E. 2020. Climate-associated genetic variation in *Fagus sylvatica* and potential responses to climate change in the French Alps. Journal of Evolutionary Biology 33: 783–796.

Capblancq T, Forester BR. 2021. Redundancy analysis: A Swiss Army Knife for landscape genomics. Methods in Ecology and Evolution 12: 2298–2309.

Catchen J, Hohenlohe PA, Bassham S, Amores A, Cresko WA. 2013. Stacks: an analysis tool set for population genomics. Molecular Ecology 22: 3124–3140.

Chase MW, Cameron KM, Freudenstein JV, Pridgeon AM, Salazar G, Van den Berg C, Schuiteman A. 2015. An updated classification of Orchidaceae. Botanical journal of the Linnean Society 177: 151–174.

Christenhusz MJ, Byng JW. 2016. The number of known plants species in the world and its annual increase. Phytotaxa 261: 201–217.

Cowling RM, Potts AJ, Bradshaw PL, Colville J, Arianoutsou M, Ferrier S, Forest F, Fyllas NM, Hopper SD, Ojeda F. 2015. Variation in plant diversity in mediterranean-climate ecosystems: The role of climatic and topographical stability. Journal of Biogeography 42: 552–564.

Cozzolino S, Widmer A. 2005. Orchid diversity: an evolutionary consequence of deception? Trends in Ecology & Evolution, 20: 487–494.

De Kort H, Vandepitte K, Bruun HH, Closset-Kopp D, Honnay O, Mergeay J. 2014. Landscape genomics and a common garden trial reveal adaptive differentiation to temperature across Europe in the tree species *Alnus glutinosa*. Molecular Ecology 23: 4709–4721.

Duffy KJ, Scopece G, Cozzolino S, Fay MF, Smith RJ, Stout JC. 2009. Ecology and genetic diversity of the dense-flowered orchid, *Neotinea maculata*, at the centre and edge of its range. Annals of Botany 104: 507–516.

Duffy KJ, Waud M, Schatz B, Petanidou T, Jacquemyn H. 2019. Latitudinal variation in mycorrhizal diversity associated with a European orchid. Journal of Biogeography 46: 968–980.

Duffy KJ, Cafasso D, Ren MX, Cozzolino S. 2020. High haplotype diversity with fine-scale structure in a recently established population of an endangered orchid. Plant Species Biology 35: 224–232.

Edgar RC, Haas BJ, Clemente JC, Quince C, Knight R. 2011. UCHIME improves sensitivity and speed of chimera detection. Bioinformatics 27: 2194–2200.

Feliner GN. 2014. Patterns and processes in plant phylogeography in the Mediterranean Basin. *A review*. Perspectives in Plant Ecology, Evolution and Systematics 16: 265–278.

Fick SE, Hijmans RJ. 2017. WorldClim, *2:* new 1-km spatial resolution climate surfaces for global land areas. International Journal of Climatology 37: 4302–4315.

Fischer MC, Rellstab C, Tedder A, Zoller S, Gugerli F, Shimizu KK, Holderegger R, Widmer A 2013. Population genomic footprints of selection and associations with climate in natural populations of *Arabidopsis halleri* from the Alps. Molecular Ecology 22: 5594–5607.

Forester BR, Lasky JR, Wagner HH, Urban DL. 2018. Comparing methods for detecting multilocus adaptation with multivariate genotype–environment associations. Molecular Ecology 27: 2215–2233.

Frei E R, Ghazoul J, Matter P, Heggli M, Pluess AR. 2012. Plant population differentiation and climate change: responses of grassland species along an elevational gradient. Global Change Biology 18: 778–789.

Frichot E, Mathieu F, Trouillon T, Bouchard G, François O. 2014. Fast and efficient estimation of individual ancestry coefficients. Genetics 196: 973–983.

Frichot E, François O. 2015. LEA: An R package for landscape and ecological association studies. Methods in Ecology and Evolution 6: 925–929.

Gallegos C, Hodgins KA, Monro K. 2023. Climate adaptation and vulnerability of foundation species in a global change hotspot. Molecular Ecology 32: 1990–2004.

Galmés J, Medrano H, Flexas J. 2007. Photosynthetic limitations in response to water stress and recovery in Mediterranean plants with different growth forms. New Phytologist 175: 81–93.

Genre A, Ivanov S, Fendrych M, Faccio A, Žárský V, Bisseling T, Bonfante P. 2012. Multiple exocytotic markers accumulate at the sites of perifungal membrane biogenesis in arbuscular mycorrhizas. Plant and Cell Physiology 53: 244–255.

Givnish TJ. 2010. Ecology of plant speciation. Taxon 59, 1326–1366.

Gugger PF, Fitz-Gibbon ST, Albarrán-Lara A, Wright JW, Sork VL. 2021. Landscape genomics of *Quercus lobata* reveals genes involved in local climate adaptation at multiple spatial scales. Molecular Ecology 30: 406–23.

Hamrick JL, Trapnell DW. 2011. Using population genetic analyses to understand seed dispersal patterns. Acta Oecologica 37: 641–649.

Hartke J, Waldvogel A, Sprenger PP, Schmitt T, Menzel F, Pfenninger M, Feldmeyer B. 2021. Little parallelism in genomic signatures of local adaptation in two sympatric, cryptic sister species. Journal of Evolutionary Biology 34: 937–952.

Hawkins BA, Field R, Cornell HV, Currie DJ, Guégan JF, Kaufman DM, Kerr JT, Mittelbach GG, Oberdorff T, O’Brien EM, Porter EE, Turner JRG. 2003. Energy, water, and broad-scale geographic patterns of species richness. Ecology 84: 3105–3117.

Hijmans RJ, Williams E, Vennes C, Hijmans MRJ. 2017. Package ‘geosphere’. Spherical trigonometry 1: 1–45.

Holderegger R, Buehler D, Gugerli F, Manel S 2010. Landscape genetics of plants. Trends in Plant Science 15: 675–683.

Hoban S, Kelley J, Lotterhos KE, Antolin MF, Bradburd G, Lowry DB, Poss ML, Reed LK, Storfer A, Whitlock MC, 2016. Finding the genomic basis of local adaptation: pitfalls, practical solutions, and future directions. The American Naturalist 188: 379–397.

Hoskin CJ, Higgie M, McDonald KR, Moritz C 2005. Reinforcement drives rapid allopatric speciation. Nature 437: 1353–1356.

Ihrmark K, Bödeker IT, Cruz-Martinez K, Friberg H, Kubartova A, Schenck J, Strid Y, Stenlid J, Brandström-Durling M, Clemmensen KE. 2012. New primers to amplify the fungal ITS2 region–evaluation by 454-sequencing of artificial and natural communities. FEMS Microbiology Ecology 82: 666–677.

Jacquemyn H, Brys R, Vandepitte K, Honnay O, Roldán-Ruiz I, Wiegand T 2007. A spatially explicit analysis of seedling recruitment in the terrestrial orchid *Orchis purpurea*. New Phytologist 176: 448–459.

Jacquemyn H, Merckx V, Brys R, Tyteca D, Cammue BP, Honnay O, Lievens B. 2011. Analysis of network architecture reveals phylogenetic constraints on mycorrhizal specificity in the genus *Orchis* (Orchidaceae). New Phytologist 192: 518–28.

Jacquemyn H, Deja A, Bailarote BC, Lievens B. 2012. Variation in mycorrhizal associations with tulasnelloid fungi among populations of five *Dactylorhiza* species. PLoS ONE, 7: e42212.

Jia K, Zhao W, Maier PA, Hu X, Jin Y, Zhou S, Jiao S, El-Kassaby YA, Wang T, Wang X, Mao J. 2020. Landscape genomics predicts climate change-related genetic offset for the widespread *Platycladus orientalis* (Cupressaceae). Evolutionary Applications 13: 665–676.

Jombart T, Devillard S, Balloux F 2010. Discriminant analysis of principal components: a new method for the analysis of genetically structured populations. BMC Genetics 11: 1–15.

Kalinowski ST. 2011. The computer program STRUCTURE does not reliably identify the main genetic clusters within species: simulations and implications for human population structure. Heredity 106: 625–632.

Kawecki TJ, Ebert D 2004. Conceptual issues in local adaptation. Ecology Letters 7: 1225–1241.

Kiers ET, Duhamel M, Beesetty Y, Mensah JA, Franken O, Verbruggen E, Fellbaum CR, Kowalchuk GA, Hart MM, Bago A. 2011. Reciprocal rewards stabilize cooperation in the mycorrhizal symbiosis. Science 333: 880–882

Kõljalg U, Nilsson RH, Abarenkov K, Tedersoo L, Taylor AF, Bahram M, Bates ST, Bruns TD, Bengtsson-Palme J, Callaghan TM. 2013. Towards a unified paradigm for sequence-based identification of fungi. Molecular Ecology 22: 5271–5277

Kretzschmar H, Eccarius W, Dietrich H. 2007. The Orchid Genera *Anacamptis, Orchis, Neotinea*: Phylogeny, Taxonomy, Morphology, Biology, Distribution. Ecology and Hybridisation. Bürgel.

Lawson DJ, Van Dorp L, Falush D. 2018. A tutorial on how not to over-interpret STRUCTURE and ADMIXTURE bar plots. Nature communications 9: 3258.

Lenormand T 2002. Gene flow and the limits to natural selection. Trends in Ecology & Evolution 17: 183–189.

Lewis ME, Puterman ML. 2001. A probabilistic analysis of bias optimality in unichain Markov decision processes. IEEE Transactions on Automatic Control 46: 96–100.

Luu, K, E Bazin, and M G B Blum. 2017. “Pcadapt: An R Package to Perform Genome Scans for Selection Based on Principal Component Analysis.” Molecular Ecology Resources 17: 67–77.

Li M-H, Liu K-W, Li Z, Lu H-C, Ye Q-L, Zhang D, Wang J-Y, Li Y-F, Zhong Z-M, Liu X. 2022. Genomes of leafy and leafless *Platanthera* orchids illuminate the evolution of mycoheterotrophy. Nature Plants 8: 373–388.

Médail F, Diadema K. 2009. Glacial refugia influence plant diversity patterns in the Mediterranean Basin. Journal of biogeography 36: 1333–1345.

McCormick MK, Jacquemyn H. 2014. What constrains the distribution of orchid populations? New Phytologist 202: 392–400.

McCormick MK, Whigham DF, Canchani-Viruet A. 2018. Mycorrhizal fungi affect orchid distribution and population dynamics. New Phytologist 219: 1207–1215.

Minasiewicz J, Znaniecka JM, Górniak M, Kawiński A. 2018. Spatial genetic structure of an endangered orchid *Cypripedium calceolus* (Orchidaceae) at a regional scale: limited gene flow in a fragmented landscape. Conservation Genetics. 19(6):1449–60.

Mosca E, Di Pierro EA, Budde KB, Neale DB, González-Martínez SC. 2018. Environmental effects on fine-scale spatial genetic structure in four Alpine keystone forest tree species. Molecular Ecology 27: 647–658.

Neiland MRM, Wilcock CC. 1998. Fruit set, nectar reward, and rarity in the Orchidaceae. American Journal of Botany 85: 1657–1671.

Nguyen LT, Schmidt HA, von Haeseler A, Minh BQ. 2015. IQ-TREE: A fast and effective stochastic algorithm for estimating maximum-likelihood phylogenies. Molecular Biology and Evolution 32: 268–274.

Oksanen J, Blanchet FG, Friendly M, Kindt R, Legendre P, McGlinn D, Minchin PR, O’hara R, Simpson GL, Solymos P 2019. Package ‘vegan’. Community ecology package. R package version 2.5-6.

Otero JT, Flanagan NS. 2006. Orchid diversity–beyond deception. Trends in Ecology & Evolution, 21: 64–65.

Parniske M. 2008. Arbuscular mycorrhiza: the mother of plant root endosymbioses. Nature Reviews Microbiology 6: 763–775.

Paz A, Ibáñez R, Lips KR, Crawford AJ. 2015. Testing the role of ecology and life history in structuring genetic variation across a landscape: a trait-based phylogeographic approach. Molecular Ecology 24: 3723–3737.

Pearson RG, Dawson TP. 2003. Predicting the impacts of climate change on the distribution of species: are bioclimate envelope models useful? Global Ecology and Biogeography 12: 361–371.

Pellegrino G, Bellusci F, Musacchio A. 2010. The effects of inflorescence size and flower position on female reproductive success in three deceptive orchids. Botanical Studies 51.

Pellegrino G, Mahmoudi M, Palermo AM. 2021. Pollen viability of Euro-Mediterranean orchids under different storage conditions: The possible effects of climate change (A Dafni, Ed.). Plant Biology 23: 140–147.

Peterson BK, Weber JN, Kay EH, Fisher HS, Hoekstra HE. 2012. Double digest RADseq: an inexpensive method for de novo SNP discovery and genotyping in model and non-model species. PLoS ONE 7: e37135.

Phillips RD, Reiter N, Peakall R 2020. Orchid conservation: from theory to practice. Annals of Botany 126: 345–362.

QGIS Development Team. 2024. QGIS Geographic Information System. Open Source Geospatial Foundation Project.

Quinlan AR. 2014. BEDTools: The Swiss-army tool for genome feature analysis. Current Protocols in Bioinformatics 47: 11121.

Ramírez-Barrera SM, Velasco JA, Orozco-Téllez TM, Vázquez-López AM, Hernández-Baños BE. 2019. What drives genetic and phenotypic divergence in the Red-crowned Ant tanager (*Habia rubica*, Aves: Cardinalidae), a polytypic species? Ecology and Evolution 9: 12339–12352.

Räsänen K, Hendry AP. 2008. Disentangling interactions between adaptive divergence and gene flow when ecology drives diversification. Ecology Letters 11: 624–636.

R Core Team. 2024. R: A language and environment for statistical computing. Vienna, Austria: R Foundation for Statistical Computing

Sexton JP, Hangartner SB, Hoffmann AA. 2014. Genetic isolation by environment or distance: which pattern of gene flow is most common? Evolution 68: 1–15.

Steinbauer MJ, Field R, Grytnes JA, Trigas P, Ah-Peng C, Attorre F, Birks HJ, Borges PA, Cardoso P, Chou CH, De Sanctis M. 2016. Topography-driven isolation, speciation and a global increase of endemism with elevation. Global Ecology and Biogeography 25:1097–1107.

Smith SE, Read DJ. 2010. Mycorrhizal symbiosis. London: Academic Press.

Taylor DL, McCormick MK. 2008. Internal transcribed spacer primers and sequences for improved characterization of basidiomycetous orchid mycorrhizas. New Phytologist 177: 1020–1033.

Tedersoo L, Bahram M, Põlme S, Kõljalg U, Yorou NS, Wijesundera R, Ruiz LV, Vasco-Palacios AM, Thu PQ, Suija A 2014. Global diversity and geography of soil fungi. Science 346: 1256688.

Těšitelová T, Jersáková J, Roy M, Kubátová B, Těšitel J, Urfus T, Trávníček P, Suda J. 2013. Ploidy-specific symbiotic interactions: divergence of mycorrhizal fungi between cytotypes of the *Gymnadenia conopsea* group (Orchidaceae). New Phytologist 199:1022–33.

Tremblay RL, Ackerman JD, Zimmerman JK, Calvo RN. 2005. Variation in sexual reproduction in orchids and its evolutionary consequences: a spasmodic journey to diversification. Biological Journal of the Linnean Society 84: 1–54.

Toczydlowski RH, Waller DM. 2019. Drift happens: Molecular genetic diversity and differentiation among populations of jewelweed (*Impatiens capensis* Meerb.) reflect fragmentation of floodplain forests. Molecular Ecology 28: 2459–2475.

Van Liedekerke M, Jones A, Panagos P 2006. ESDBv2 raster library—a set of rasters derived from the European Soil Database Distribution v2. 0. European Commission and the European Soil Bureau Network, CDROM, EUR 19945.

de Villemereuil P, Frichot É, Bazin É, François O, Gaggiotti OE. 2014. Genome scan methods against more complex models: when and how much should we trust them? Molecular Ecology 23: 2006–2019.

Vogt-Schilb H, Těšitelová T, Kotilínek M, Sucháček P, Kohout P, Jersáková J 2020. Altered rhizoctonia assemblages in grasslands on ex-arable land support germination of mycorrhizal generalist, not specialist orchids. New Phytologist 227: 1200–1212.

Wang IJ, Bradburd GS. 2014. Isolation by environment. Molecular Ecology 23: 5649–5662.

Whitlock MC, KE Lotterhos. 2015. Reliable Detection of Loci Responsible for Local Adaptation: Inference of a Null Model Through Trimming the Distribution of FST. American Naturalist 186: S24–S36.

Wright S. 1943. Isolation by distance. Genetics 28: 114–138.

Wu Z, Li X, Xie D, Wang H, Zhang Z, Xu X, Li T. 2020. Spatial genetic structuring in a widespread wetland plant on a plateau: Effects of elevation-driven geographic isolation and environmental heterogeneity. Freshwater Biology 65:1596–607.

