## Supplementary figures and images for "Interplay between abiotic conditions and mycorrhizal abundance determines differentiation and potential adaptation in a Mediterranean orchid"

### Fig.S1

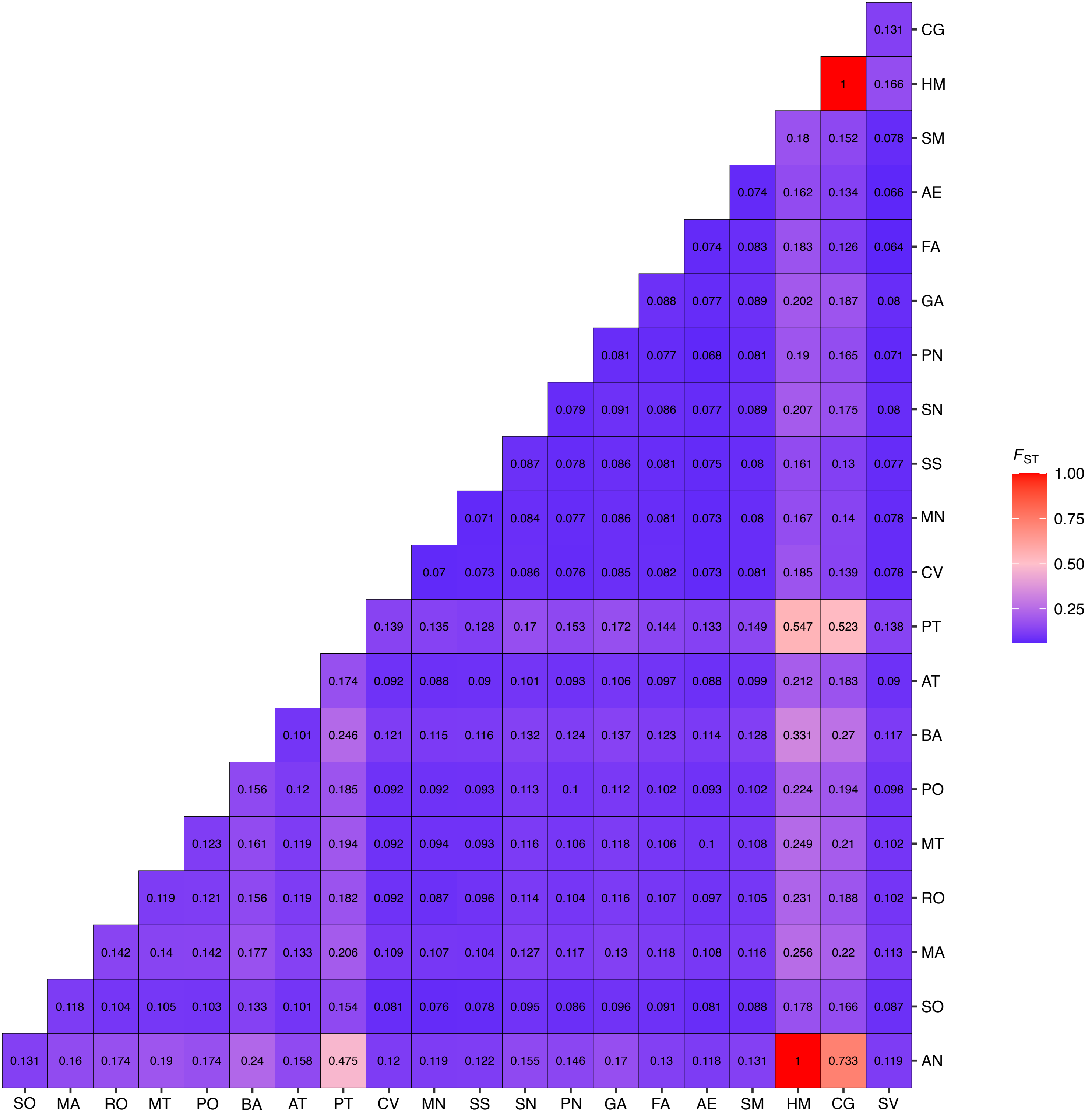

### Fig.S2

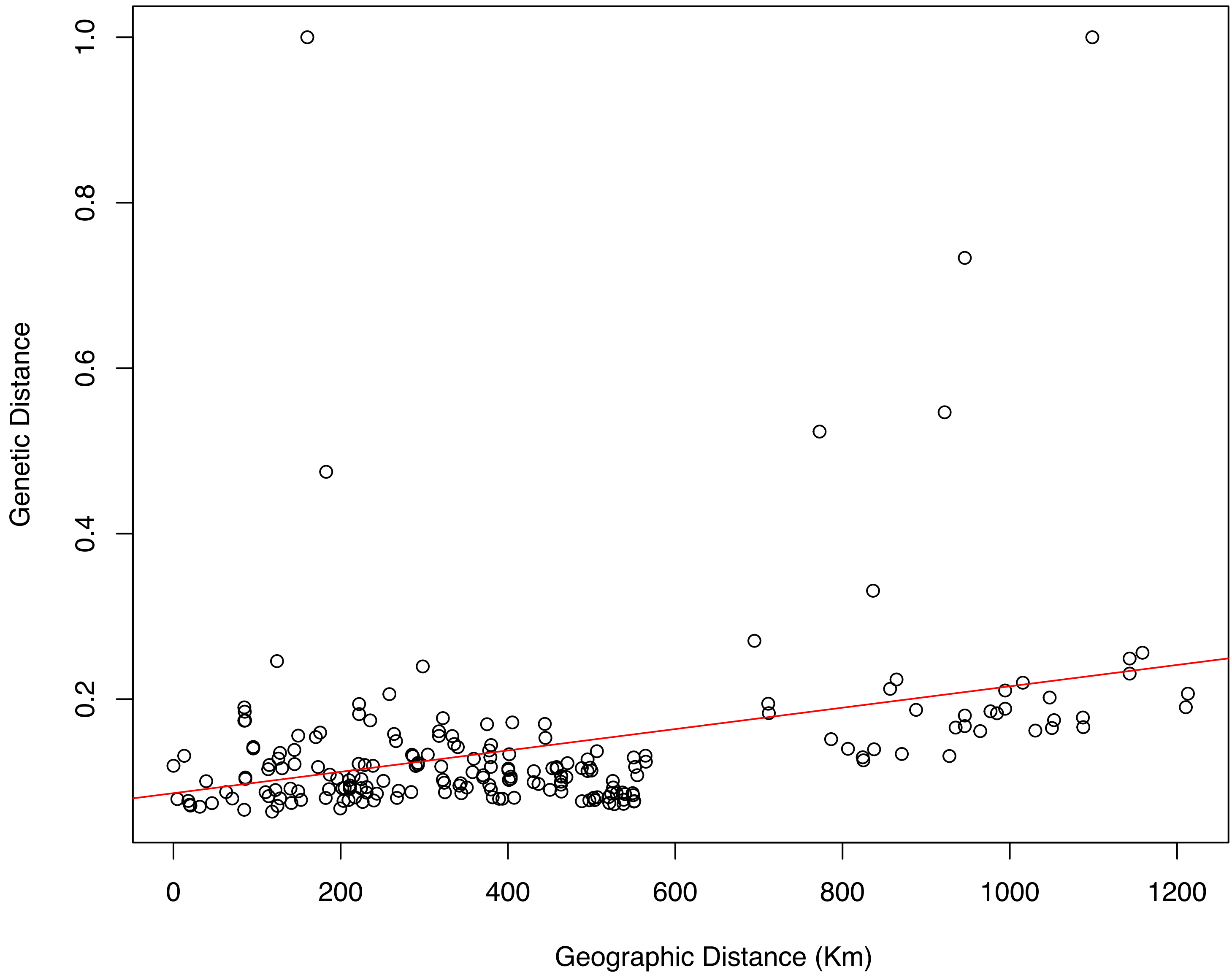

### Fig.S3

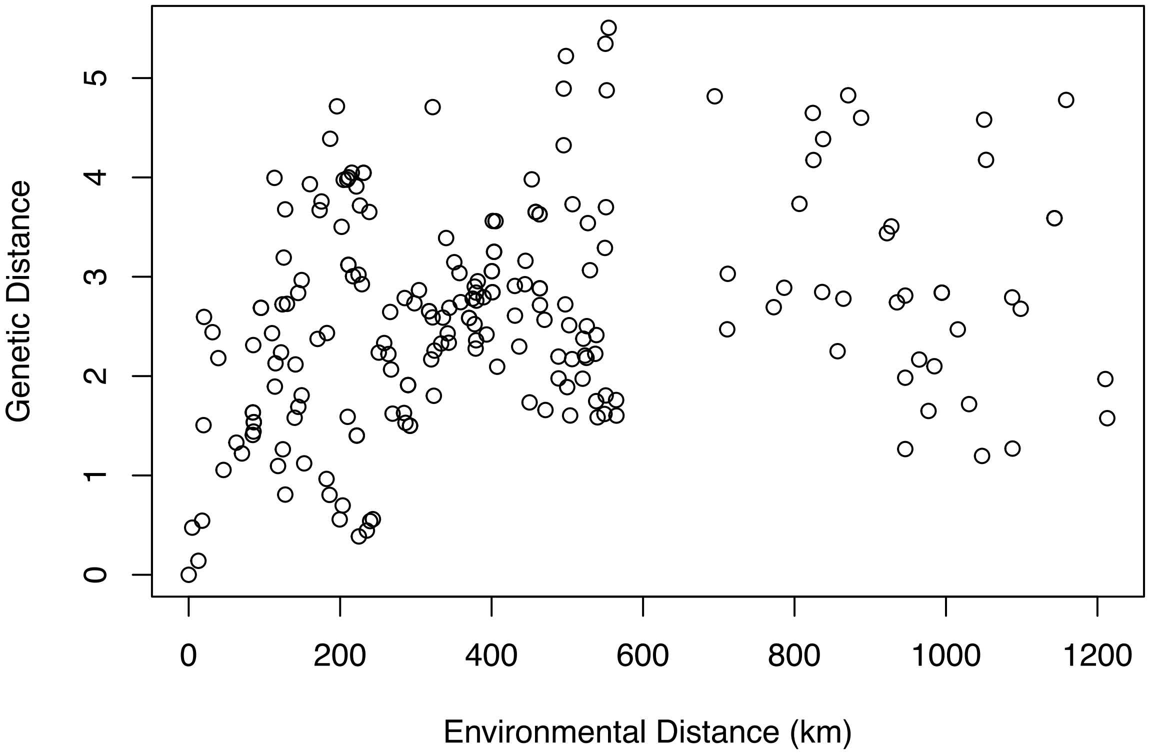

### Fig.S4

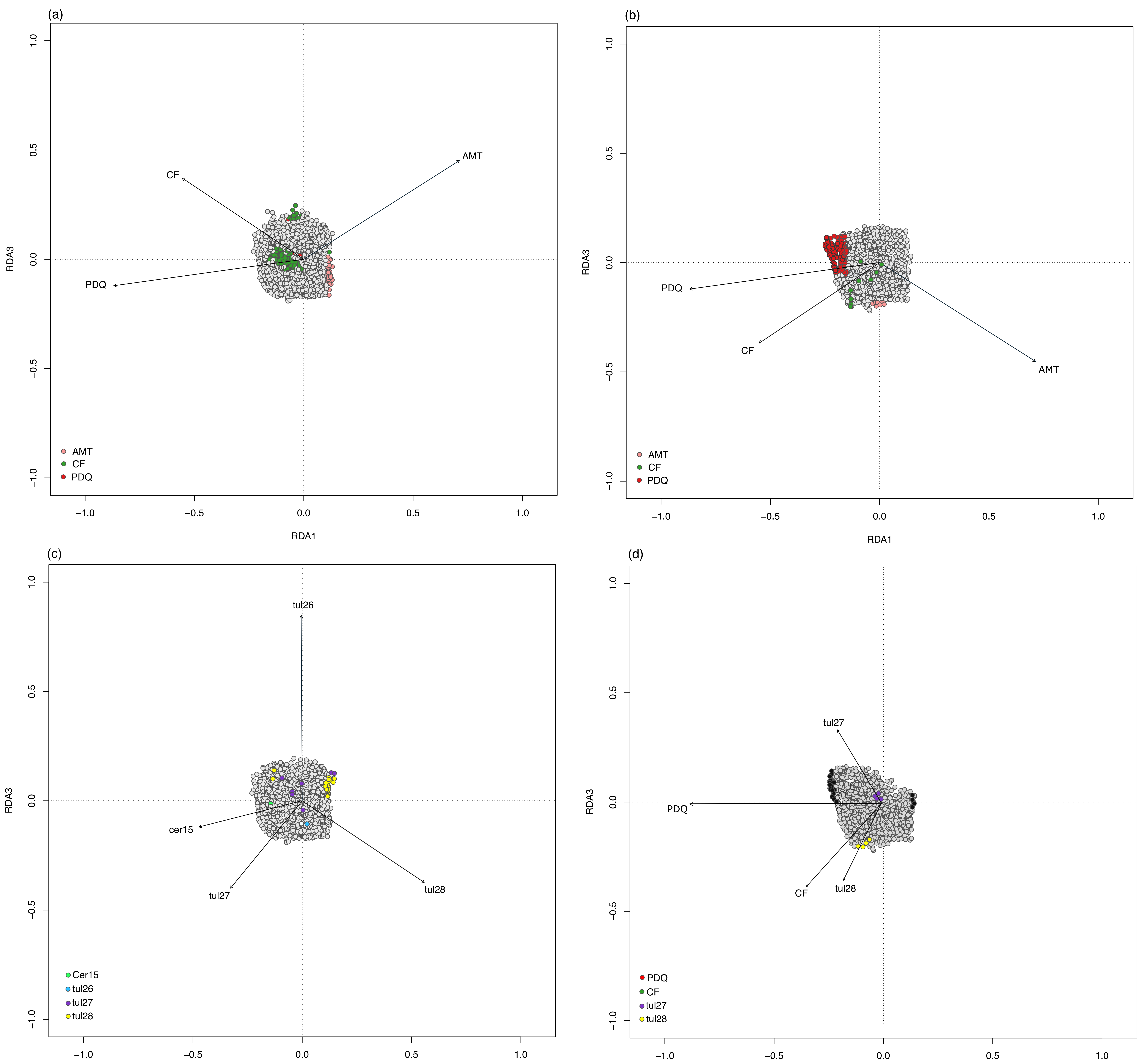

### Fig.S5

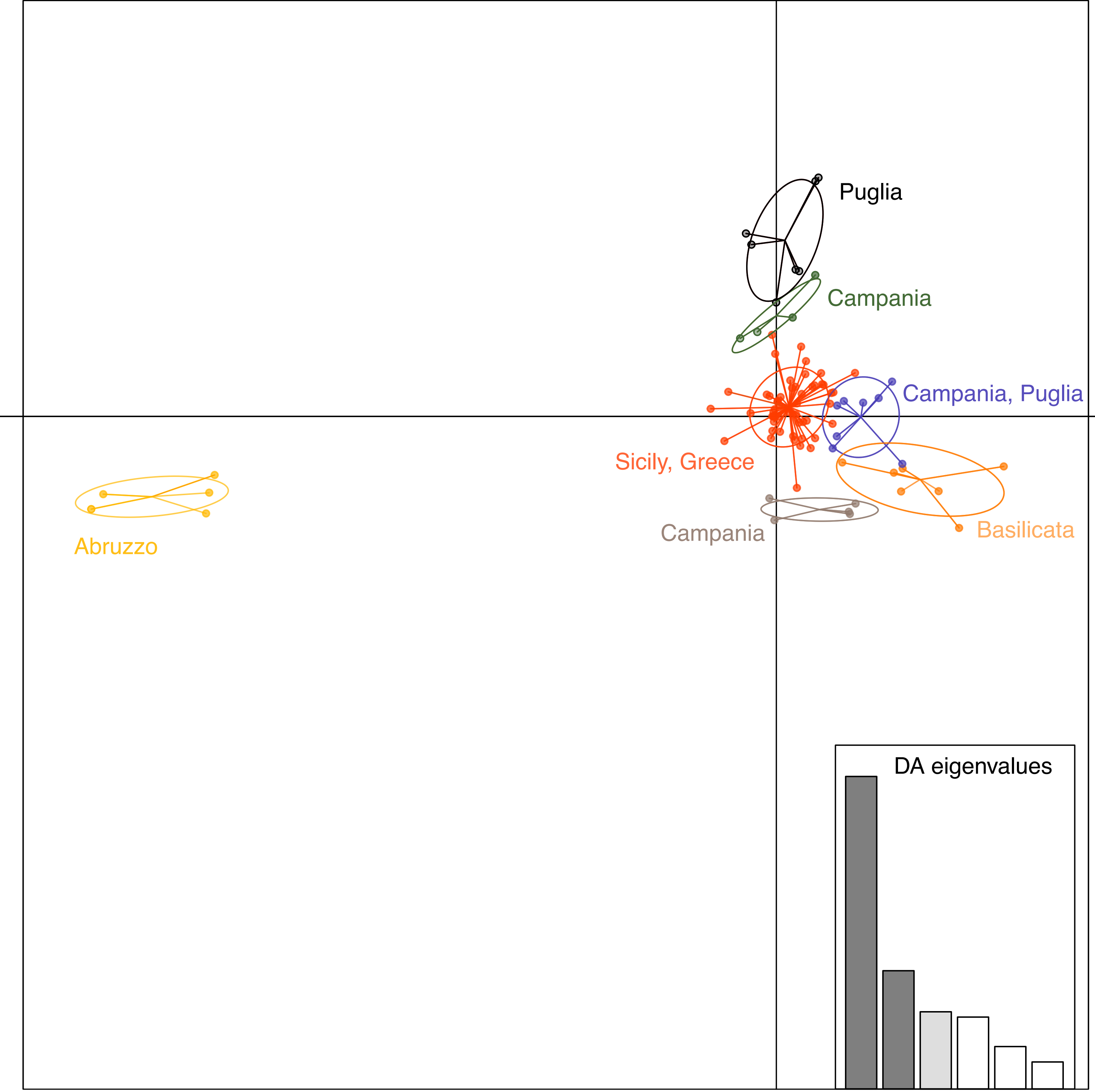

### Fig.S6

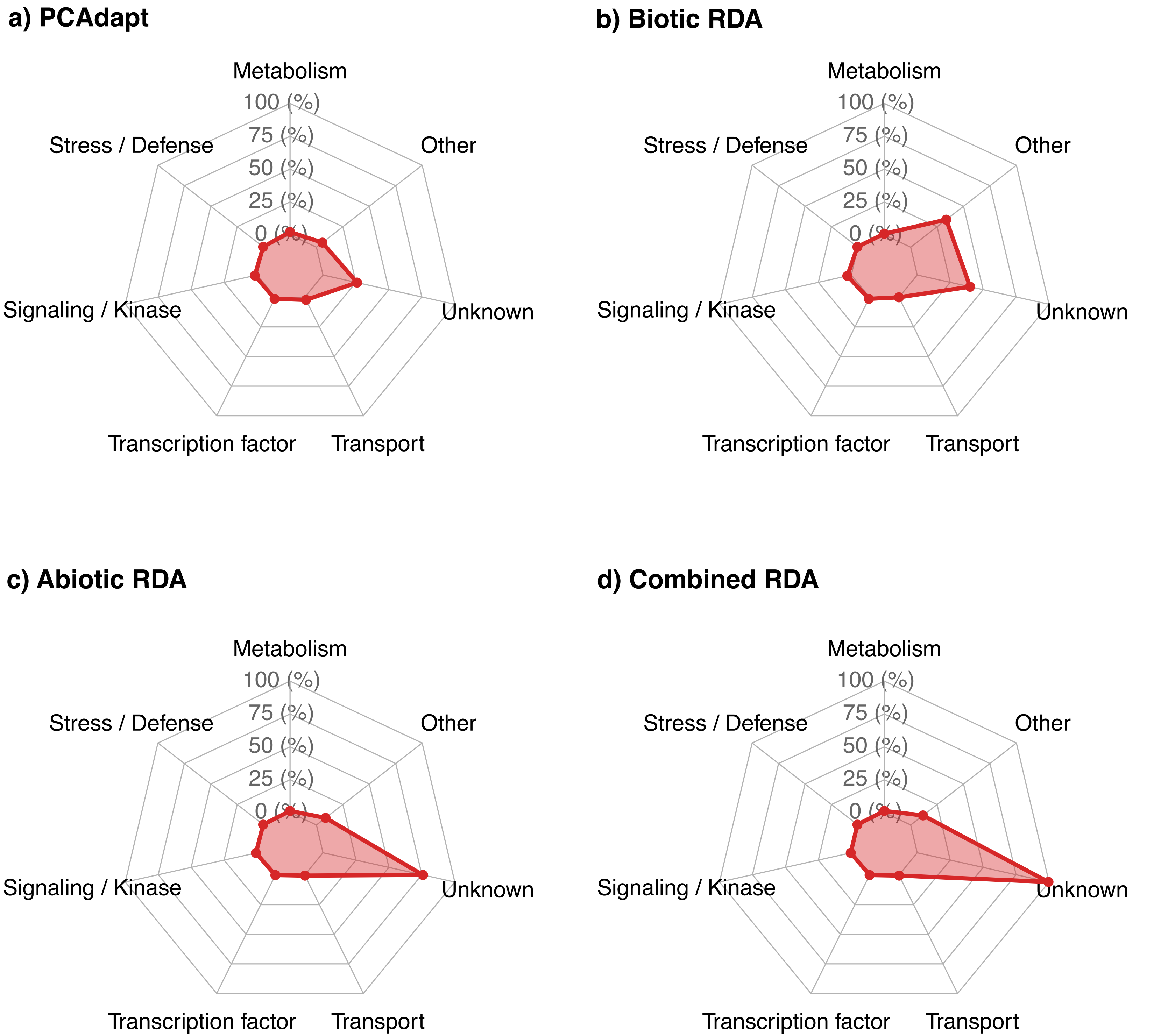
